# The gendered self: Evidence for differences in whole-brain dynamics

**DOI:** 10.1101/2021.11.19.469282

**Authors:** Carme Uribe, Anira Escrichs, Eleonora de Filippi, Yonatan Sanz-Perl, Carme Junque, Esther Gomez-Gil, Morten L Kringelbach, Antonio Guillamon, Gustavo Deco

## Abstract

How the brain constructs gender identity is largely unknown, but some neural differences have recently been discovered. Here, we used an intrinsic-ignition framework to investigate if gender identity changes the propagation of the neural activity across the whole-brain network and within resting-state networks. Studying 29 transmen and 17 transwomen with gender incongruence, 22 ciswomen, and 19 cismen, we computed the capability of a given brain area in space to propagate activity to other areas (mean-ignition) and its variability across time (node-metastability). We found that both measures differentiated all four groups across the whole-brain network. Furthermore, at the network level, we found that compared to the other groups, cismen showed higher mean-ignition of the dorsal attention network and node-metastability of the dorsal and ventral attention, executive control, and temporal parietal networks. We also found mean-ignition differences between cismen and ciswomen within the executive control network, but higher in ciswomen than cismen and transmen for the default-mode network. For the node-metastability, this was higher in cismen compared to ciswomen in the somatomotor network, while both mean-ignition and node-metastability were higher for cismen than transmen in the limbic network. Finally, we computed correlations between both measures and their body image scores. Transmen dissatisfaction, cismen, and ciswomen satisfaction towards their own body image were distinctively associated with specific networks per group. Overall, the study of the whole-brain network dynamical complexity discriminates binary gender identity groups, and functional connectivity dynamics approaches are needed to disentangle the complex understanding of the gendered self.

**Significance statement:** The study of sex/gender differences may be enriched by the heterogeneity of other gender minority groups, such as transgender. Functional connectivity measures capturing the spatio-temporal oscillations of the brain can provide insights on how the brain cooperates. This is the first study investigating how the whole-brain network propagates information across the brain, spatially and temporally, in binary gender groups (cisgender and transgender) by means of the intrinsic-ignition framework. We found four whole-brain unique phenotypes pertaining to each gender group, namely cismen, ciswomen, transmen and transwomen. Novel functional connectivity dynamics frameworks can contribute to disentangle the complex experience of a *gendered*-self.

## Introduction

A significant number of studies have explored sex-related differences in brain connectivity (Biswal et al., 2010; Ritchie et al., 2018; de Lacy et al., 2019; Eliot et al., 2021). However, the proposed sexual dimorphism, as observed in the reproductive organs, has been rejected in terms of psychological sex differences in the brain (Hyde et al., 2019; Eliot et al., 2021) and a meta-analysis informed about probable excessive significance reports, i.e., inducing a positive reporting bias (David et al., 2018). Furthermore, when investigating the brain differences between *females* and *males*, the heterogeneity of the sex/gender construct and its less prevalent forms, such as the transgender groups, have usually been overlooked. Gender identity can be defined as a complex multifactorial trait that may or may not be binary and that may correspond either to one’s sex assigned at birth, i.e., cisgender, or to a discrepant type, i.e., transgender (Polderman et al., 2018).

Understanding gender incongruence in transgender people has been a growing focus of interest, although it is important to note that not all transgender people present such incongruence. Recently, intrinsic brain functional connectivity have differentiated gender groups (Nota et al., 2017; Clemens et al., 2020; Uribe et al., 2020b). However, as the brain networks are constantly interacting (Menon, 2011; Chen et al., 2013), the understanding of their interplay across the whole-brain underlying the complex construction of gender (Uribe et al., 2020b) is something worth exploring.

The study of the brain network interactions is enriched by investigating its spatio-temporal fluctuations in response to internal and external stimuli. Whole-brain dynamics differences between cismen and ciswomen have been described using a sliding window approach (de Lacy et al., 2019), and more recently, with a small dataset of transmen (Uribe et al., 2021). This latter study mainly differentiated cismen from transmen and ciswomen, while these last groups had statistically equivalent fluidity and range dynamism (Uribe et al., 2021). On the other hand, the brain dynamics of transwomen remain elusive. Although differences in the interactions among large-scale networks between cis- and transgender groups have been described (Uribe et al., 2020b), it remains unclear how such networks cooperate to build gender identity.

In recent years, a growing number of data-driven approaches have been proposed to describe the spatiotemporal brain dynamics (Allen et al., 2014; Hansen et al., 2015; Deco et al., 2017a). Among them, the novel intrinsic ignition framework has been lately developed to investigate the propagation of activity over time across the whole-brain (Deco and Kringelbach, 2017). This data-driven method was conceived to capture the influence of local activity on the global brain computation by describing the broadness of communication (Deco and Kringelbach, 2017). In particular, the concept of intrinsic ignition reflects a degree of global integration induced by the capability of a given brain area to propagate neural activity across the whole-brain network.

In this work, we explored gender-related differences in whole-brain functional dynamics. Specifically, we investigated the dynamical complexity of four gender groups (transmen and transwomen with gender incongruence, cismen, and ciswomen) by looking at the effects of spontaneously occurring local activation (i.e., events) on global integration (Deco et al., 2015) through the intrinsic ignition framework (Deco and Kringelbach, 2017; Deco et al., 2017b). While large samples are needed to capture gender effects in the brain (Ritchie et al., 2018), we studied a well-characterized group of cisgender and transgender people beyond the assumed self-identification to a cisgender identity.

Second, we were also interested in exploring the associations of the functional connectivity dynamics with the degree of satisfaction towards body parts. Although it may not be the case for all transgender people, they frequently report gender nonconformity towards the sex assigned at birth, especially when they still have not gone through a gender-affirmative hormone treatment (GAHT) (Selvaggi and Bellringer, 2011). The transgender participants of this study presented with such disconformity, and they were candidates to initiate GAHT.

## Methods and materials

### Participants and instruments

Twenty-nine transmen with no GAHT initiated participants (age: mean(SD) = 24.7(6.2), range = 17 — 39; education: mean(SD) = 11.7(1.7), range = 9 — 15), 17 transwomen with no GAHT (age: mean(SD) = 21.4(3.9), range = 18 — 34, education: mean(SD) = 13.1(1.8), range = 10 — 16), 19 cismen (age: mean(SD) = 22.2(4.4), range = 18 — 32, education mean(SD) = 14.4(3.0), range = 10 — 20), and 22 ciswomen (age: mean(SD) = 19.6(2.4), range = 18 — 27, education: mean(SD) = 13.3(1.6), range = 12 — 17) were enrolled. All participants explicitly stated to have a binary identity, thus transmen and cismen identified themselves as a man, and transwomen and ciswomen as a woman. Detailed demographic information such as age and education, and information of the protocol assessment and recruitment can be found in the data article (Uribe et al., 2020a). All transmen and transwomen met diagnostic criteria for gender identity disorder according to the DSM-IV-TR and ICD-10 when recruited. Nonetheless, the diagnosis was relabeled to gender incongruence as per the ICD-11 and recommended by EPATH and WPATH v7 (Bouman et al., 2017).

Participants answered the Body Image Scale (Lindgren and Pauly, 1975). This auto-administered questionnaire has a total averaged score that includes: the nose, shoulders, chin, calves, hands, adam’s apple, eyebrows, face, feet, height, hips, figure, waist, arms, buttocks, biceps, appearance, stature, muscles, weight, thighs, breasts, chest, body hair, facial hair, hair, voice, penis/vagina, scrotum/clitoris, and testicles/uterus. Participants scored each body part on a Likert scale ranging from 1 (Very satisfied) – 2 (Satisfied) – 3 (Neutral) – 4 (Dissatisfied) – 5 (Very dissatisfied). Written informed consent was obtained from all participants after a full explanation of the procedures. The study was approved by the ethics committee of the Hospital Clinic of Barcelona.

### MRI acquisition and preprocessing

Raw and processed imaging data are available online (Uribe et al., 2020a). MRI data were acquired with a 3T scanner (MAGNETOM Trio, Siemens, Germany). Briefly, T1-weighted images were acquired in the sagittal plane, TR = 2,300 ms, TE = 2.98 ms, TI = 900 ms, 240 slices, FOV = 256 mm; matrix size = 256 × 256; 1 mm isotropic voxel. A total of 240 T2* weighted images were acquired with a TR = 2,500 ms s, TE = 28 ms, flip angle = 80°, slice thickness = 3 mm, FOV = 240 mm. Participants were instructed not to fall asleep and not to focus in any specific thought, keeping their eyes closed. Basic preprocessing was conducted with AFNI using an in-house shell script. ICA-AROMA was applied for the automatic removal of motion-related artifact. No motion parameter differed between groups (Uribe et al., 2020a).

We extracted the timeseries of the 1,000 nodes parcellation in Schaefer et al. (2018) from Yeo’s resting-state networks (Thomas Yeo et al., 2011) with the *fslmeants* tool from FSL v5.0.10 (https://fsl.fmrib.ox.ac.uk/fsl/fslwiki/). The 17-network Schaefer parcellation was used to define networks by grouping, for example, the A, B, and C components of the default-mode or the executive control networks as one. We chose the 17 partition as there was the unique temporal parietal network, otherwise subdivided by several other networks in the 7-network partition, namely the default-mode, ventral attention, and somatomotor networks. For a comprehensive description of the method we refer readers to Schaefer et al. (2018) and Thomas Yeo et al. (2011).

### Intrinsic ignition framework

We applied the intrinsic ignition framework to characterize gender-related differences in the spatiotemporal transmission of information across the entire brain over time (Deco and Kringelbach, 2017; Deco et al., 2017b). This framework has been used to successfully discriminate between different brain states, such as sleep (Deco et al., 2017b) and meditation (Escrichs et al., 2019), to explore differences in the healthy elderly brain (Escrichs et al., 2020), in depression (Alonso Martínez et al., 2020; Mayneris-Perxachs et al., 2021), and even in preterm children (Padilla et al., 2020). It allows computing the effect of spontaneous local activation on the whole-brain integration using the phase space of the signals. First, we filtered the blood-oxygen-level-dependent (BOLD) time-series parcellated in 1,000 brain regions within the narrowband 0.04 — 0.07 Hz to avoid artifacts (Glerean et al., 2012) (Figure 1.1A). Then, we calculated the instantaneous phase of the BOLD signals by computing the Hilbert Transform of the filtered time-series (Figure 1.1B).

**Figure 1:**
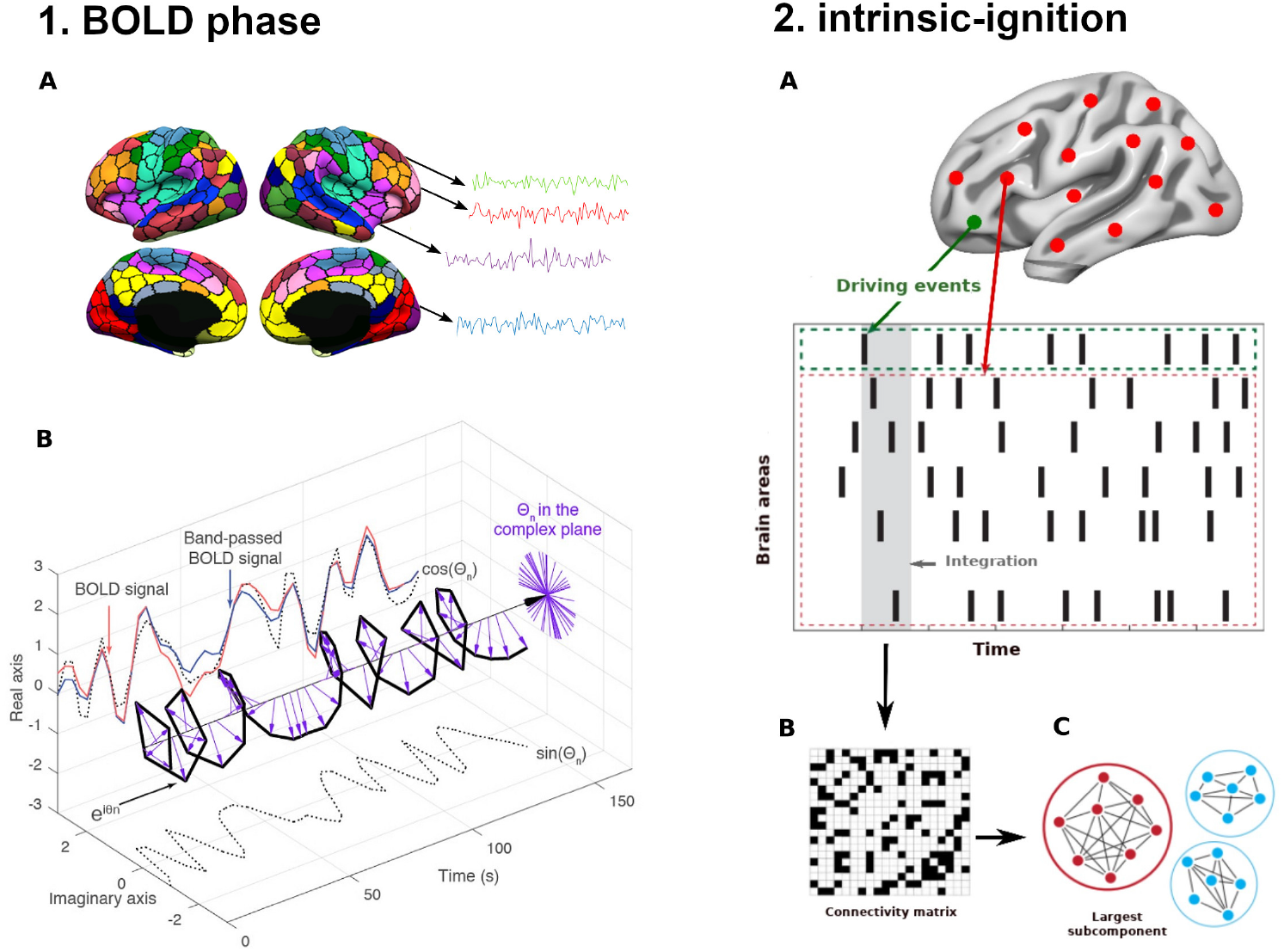
Intrinsic ignition framework. **(1)** We extracted the BOLD time series for each of the 1,000 brain areas and computed the phase space of the BOLD signal. **(1A)** We obtained the time series of each parcellation using the resting-state Schaefer atlas (Schaefer et al., 2018). **(1B)** Then, we measured the phase space of the BOLD signal through the Hilbert transform for each region. The BOLD signal (red) was band-pass filtered between 0.04 — 0.07 Hz (blue) and using the Hilbert transform. The phase dynamics can be represented in the complex plane as e^*iϕ*^ (black bold line), the real part as cos *ϕ* and the imaginary part as sin *ϕ* (black dotted lines). The purple arrows represent the Hilbert phases at each TR (2.5s). **(2)** Intrinsic ignition measurements. **(2A)** Events were captured applying a threshold method (Tagliazucchi et al., 2012) (see green node). For each event elicited, the activity in the rest of the network (see red stippled region) was measured in the time-window of 4-TR (4×2.5s) (gray area). **(2B)** A binarized phase lock matrix was obtained from the time-window. **(2C)** From this phase-lock matrix, we obtained the integration measure by computing the largest subcomponent, i.e., by applying the global integration measure (Deco et al., 2015, 2017b). Repeating the procedure for each driving event, we obtained the ignition and node-metastability of the intrinsic-driven integration for each brain region across the whole-brain network. Figure adapted from (Deco and Kringelbach, 2017; Deco et al., 2019; Escrichs et al., 2020).

Figure 1.2 gives a graphical representation of the algorithm used to compute the ignition value for each brain area evoked for an event within a fixed time window of 4-TR (TR = 2.5s). In brief, a binary event was defined by transforming the time series into z-scores, *z_i_*(t), and by fixing a threshold *θ* (Tagliazucchi et al., 2012; Deco et al., 2017b). Then, the phase lock matrix *P_jk_*(*t*) was calculated, representing the state of pair-wise phase synchronization at each time-point *t* between regions *j* and *k*, as given by:

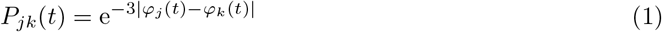

where *φ_j_*(*t*) and *φ_k_*(*t*) represent the obtained phase at time *t* of the regions *j* and *k*. Given the fixed threshold *θ*, the symmetric phase lock matrix *P_jk_*(*t*) was binarized (Figure 1.2B) such that *σ*(t)=1 if *z_i_*(t) > *θ* and 0 otherwise. We computed the integration value as the length of the connected component (i.e., the largest subcomponent) considered as an adjacent graph (Figure 1.2C). Finally, we obtained the average integration value (i.e., ignition) by averaging across all events and the variability (i.e., node-metastability) by calculating the standard deviation, reflecting the spatial diversity across the whole-brain network and the level of variability over time for each brain region, respectively. The framework was applied to the whole-brain network parcellated into 1,000 brain areas and to each resting-state network separately: the dorsal and ventral attentional, executive control, default-mode, somatomotor, limbic, visual, and temporal parietal networks (Schaefer et al., 2018; Thomas Yeo et al., 2011).

**Figure 2:**
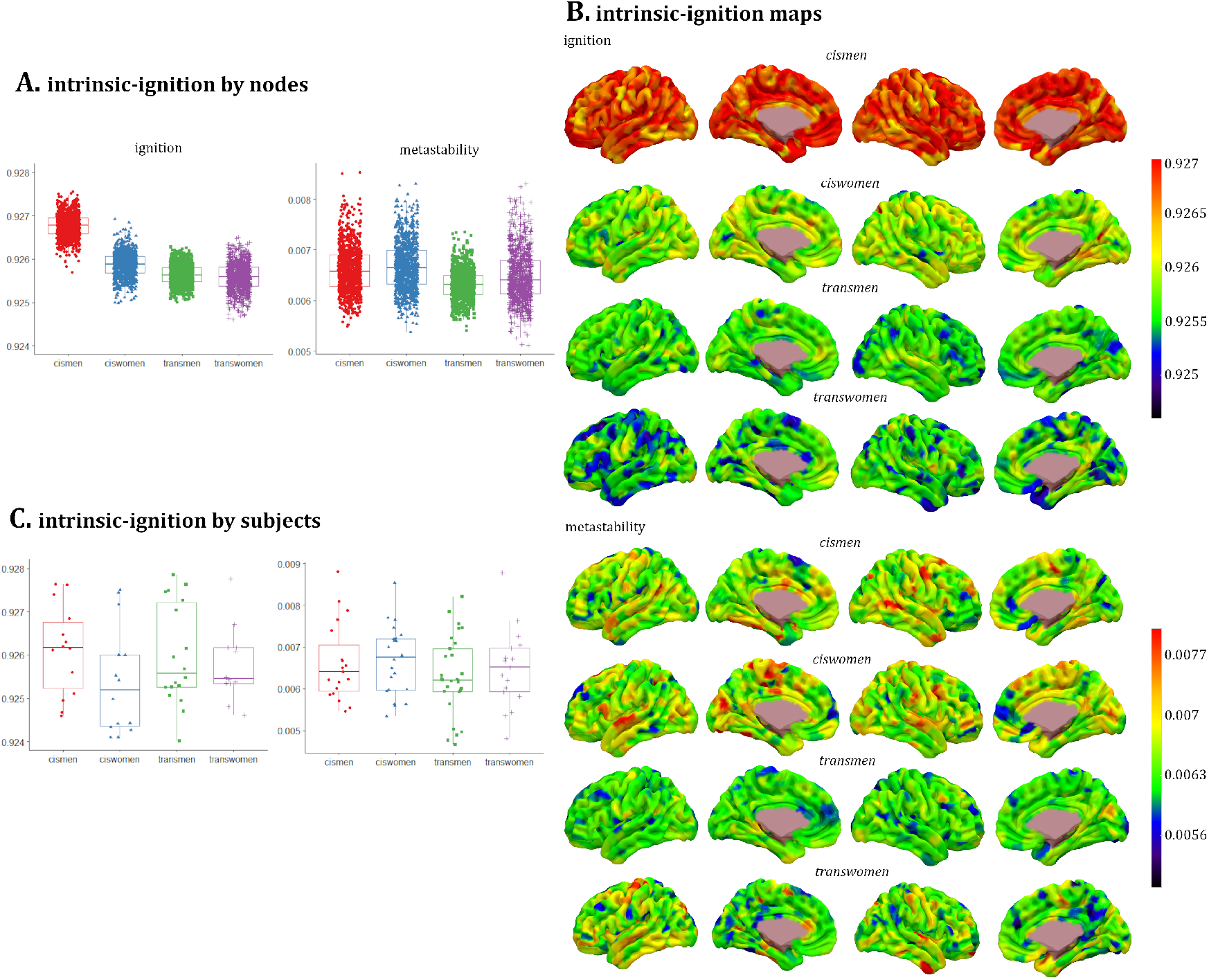
Whole-brain ignition and node-metastability measures for each of the 1,000 brain regions by each group. **(A)** The boxes in the plots indicate the second and third quartile (IQR), and middle lines are medians. Each dot represents a brain region. Means and standard deviations can be found in Table 1. There were significant differences between all groups’ contrasts with Monte-Carlo 1,000 permutations and Bonferroni correction *p* = 0.0004. **(B)** Rendered brains show the distribution of ignition and node-metastability values per each brain region by group. Red warm regions had the highest ignition and node-metastability values, and dark blue ones the lowest. **(C)** Ignition and node-metastability measures with averaged regions for each participant by groups. Each dot represents a participant. No group comparison reached the significance threshold after Bonferroni correction. Metadata can be downloaded at https://doi.org/10.6084/m9.figshare.14622564.

**Table 1:**
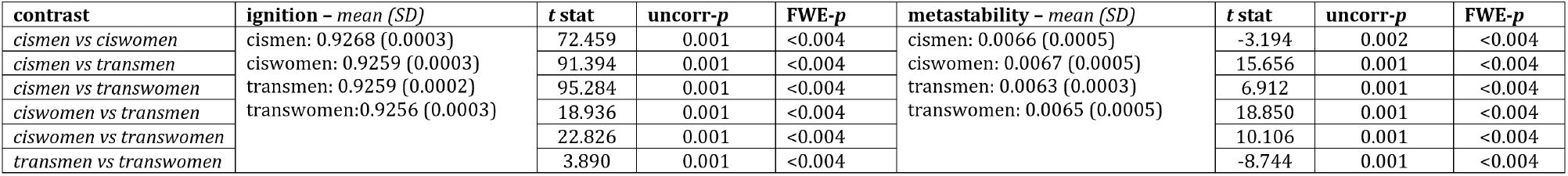
Stats from whole-brain intrinsic ignition framework by nodes. Data are means and standard deviations. There were significant differences between all contrasts groups with Monte-Carlo 1,000 permutations and family-wise error (FWE) correction.

### Statistical analyses

A general linear model and Monte Carlo permutation testing (1,000 iterations) to control for Family-Wise Error (FWE) rate were applied to perform group comparisons. Cohens’s *d* effect sizes were also calculated. Age and education were entered as covariates when comparing transmen and ciswomen, and the education variable alone when comparing transmen and cismen. Spearman correlations and 95% confidence interval were computed between the quantitative functional connectivity dynamics metrics and the body image scale total score per group. We did not include the visual network in the correlations, as there were no group differences within this network in ignition or node-metastability.

## Results

### Ignition across nodes

There were significant differences (FWE corrected *p* = 0.0042) between all gender groups when computing the intrinsic ignition framework across the whole-brain functional network in both mean-ignition and node-metastability (Figure 2A and Table 1). There was a gradual progression in the mean-ignition, i.e., cismen>ciswomen>transmen>transwomen. On the other hand, the average node-metastability was the highest in the ciswomen group, and the lowest average was observed in the transmen group.

Figure 2B shows rendered brains, where the hot colors represent those regions with the highest ignition and node-metastability per group, while cold and dark tonalities represent the lowest values. The regional distribution of the highest ignition and node-metastability measures included regions across the whole-brain from all networks. There was no hemisphere predominance among the 100 areas with the highest ignition in any group (left/right: cismen 54/46 —out of the 100 regions—, ciswomen 43/57, transmen 51/49, transwomen 38/62, *χ*^2^=6.472; *p* = 0.091). Nodes in the right hemisphere were more frequent for the node-metastability measures except in the transwomen group (left/right: cismen 41/59, ciswomen 47/53, transmen 48/52, transwomen 61/39, *χ*^2^=8.512; *p* = 0.037).

### Ignition across participants

**Whole-brain.** There were no group differences that survived FWE correction (Figure 2C). When computing the intrinsic ignition framework by networks, group differences were present in all networks except for the visual network (Table 2, Figures 3 and 4). **Attentional network**. In the dorsal attention network, the ignition (Figure 3.1C and 3.2C) and node-metastability (Figure 4.1B and 4.2B) measurements were higher in the cismen’s group with respect to ciswomen, transmen, and transwomen. Regarding the ventral subdivision of the attentional network, only the node metastability mean of cismen was higher than the other three gender groups, namely ciswomen, transmen, and transwomen (Figure 4.1C and 4.2C). **Executive control network**. Cismen’s ignition was significantly higher than ciswomen (Figure 3.1A and 3.2A). Cismen also had higher node-metastability than ciswomen, transmen, and transwomen (Figure 4.1A and 4.2A). **Default-mode network**. Ciswomen had higher ignition than cismen and transmen (Figure 3.1B and 3.2B). **Limbic network.** Both ignition and node-metastability measurements were higher within the cismen group with respect to transmen (Figures 3.1D, 3.2D, and 4.1D and 4.2D). **Somatomotor.** Cismen showed higher node-metastability than ciswomen (Figure 4.1E and 4.2E). **Temporal parietal.** Cismen had greater mean igniton than ciswomen, transmen, and transwomen (Figure 3.1E and 3.2E).

**Table 2:**
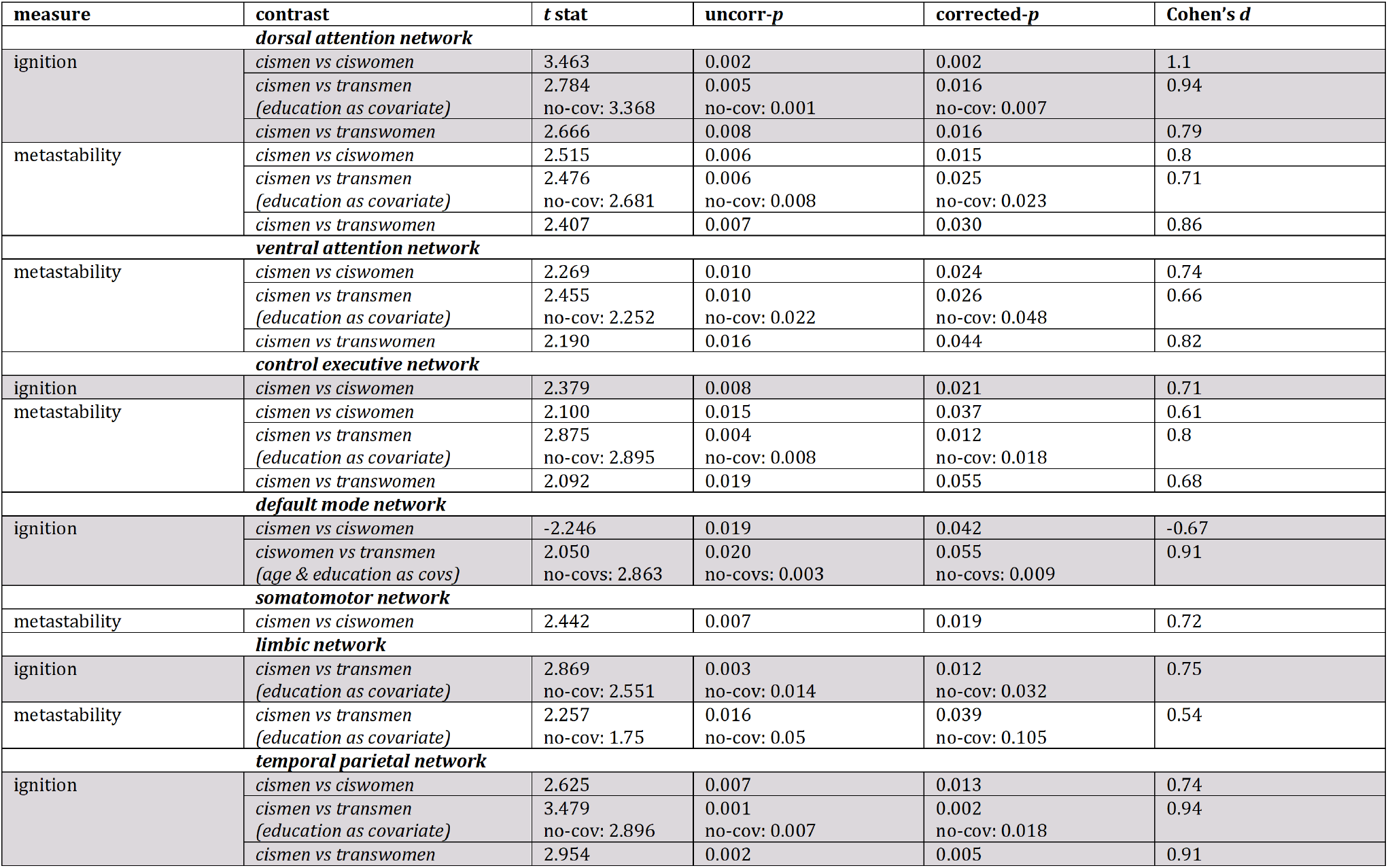
Intrinsic ignition by networks of averaged nodes across groups. General linear model with 1,000 permutations and family-wise error correction were applied. Cohen’s d effect sizes were computed. The cismen vs. transmen contrast was tested with and without (no-cov) education as covariate, and the comparison between ciswomen and transmen with and without (no-covs) age and years of education.

**Figure 3:**
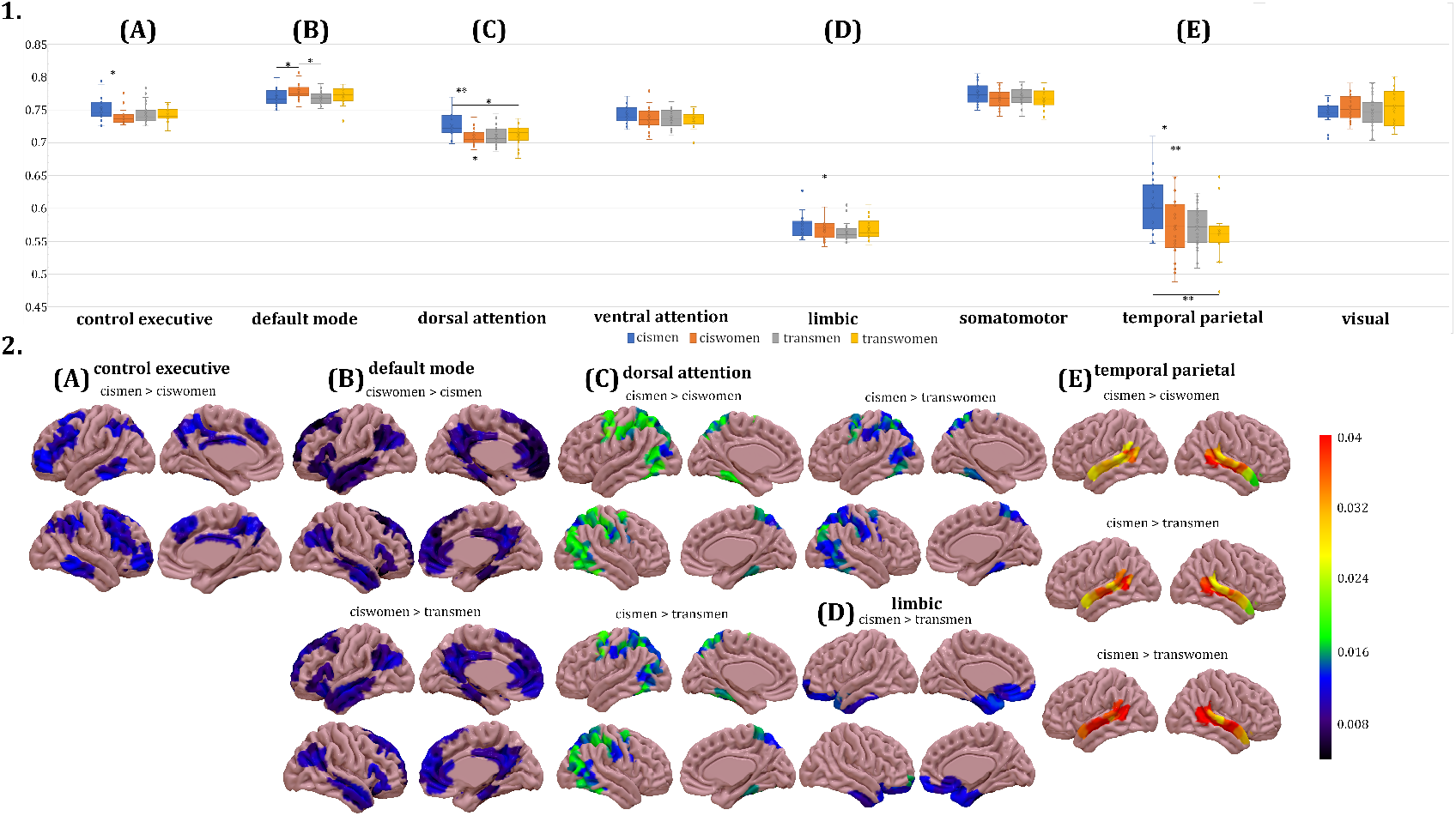
Ignition measures by networks. **(1)** Group ignition values by networks. The boxes in the plots indicate the second and third quartile (IQR), middle lines are ignition medians, and X are ignition means. Legend: * *p ≤* 0.05; ** *p* < 0.01. **(2)** Rendered brains represent the differences in ignition between groups, and were plotted with the SurfIce software. There were group differences in the **(A)** executive control, **(B)** default-mode, **(C)** dorsal attentional, **(D)** limbic, and **(E)** temporal parietal networks. Metadata can be downloaded at https://doi.org/10.6084/m9.figshare.14622564.

**Figure 4:**
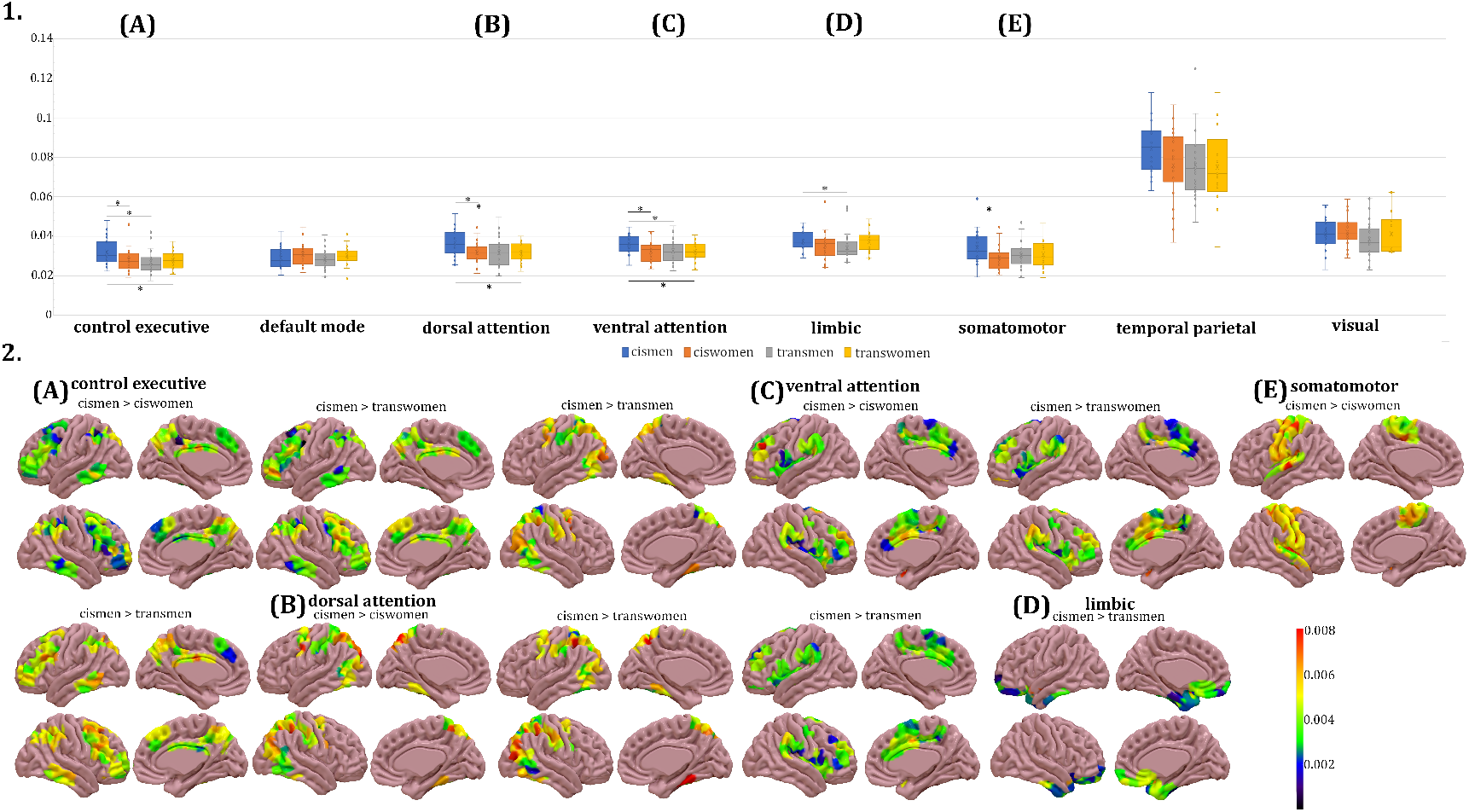
Node-metastability measures by networks. **(1)** Group node-metastability values by networks. The boxes in the plots indicate the second and third quartile (IQR), middle lines are ignition medians, and X are node-metastability means. Legend: * *p ≤* 0.05; ** *p* < 0.01. **(2)** Rendered brains depict the node-metastability difference between groups, and were plotted with the SurfIce software. There were group differences in the **(A)** executive control, **(B)** dorsal attention, **(C)** ventral attention, **(D)** limbic, and **(E)** somatomotor networks. Metadata can be downloaded at https://doi.org/10.6084/m9.figshare.14622564.

### Body image satisfaction correlations

The degree of dissatisfaction towards body parts was significantly higher in the two transgender groups in comparison to both cismen and ciswomen groups. On the other hand, satisfaction in the two cisgender groups was not statistically different (Figure 5A). The satisfaction/dissatisfaction with the body image within each gender group was associated distinctively with intrinsic ignition measures in specific networks. Cismen were the group with the highest overall mean satisfaction with their body, and this correlated negatively with the ignition in the executive control network and positively with node-metastability in the limbic network (Figure 5B). On the other hand, body image satisfaction in ciswomen correlated with the ignition in the default-mode network, which was reported to be higher in the ciswomen groups comparisons, and the temporal parietal’s node-metastability (Figure 5C). Ventral attentional ignition was positively associated with the global score of the body image scale in transmen (Figure 5D).

**Figure 5:**
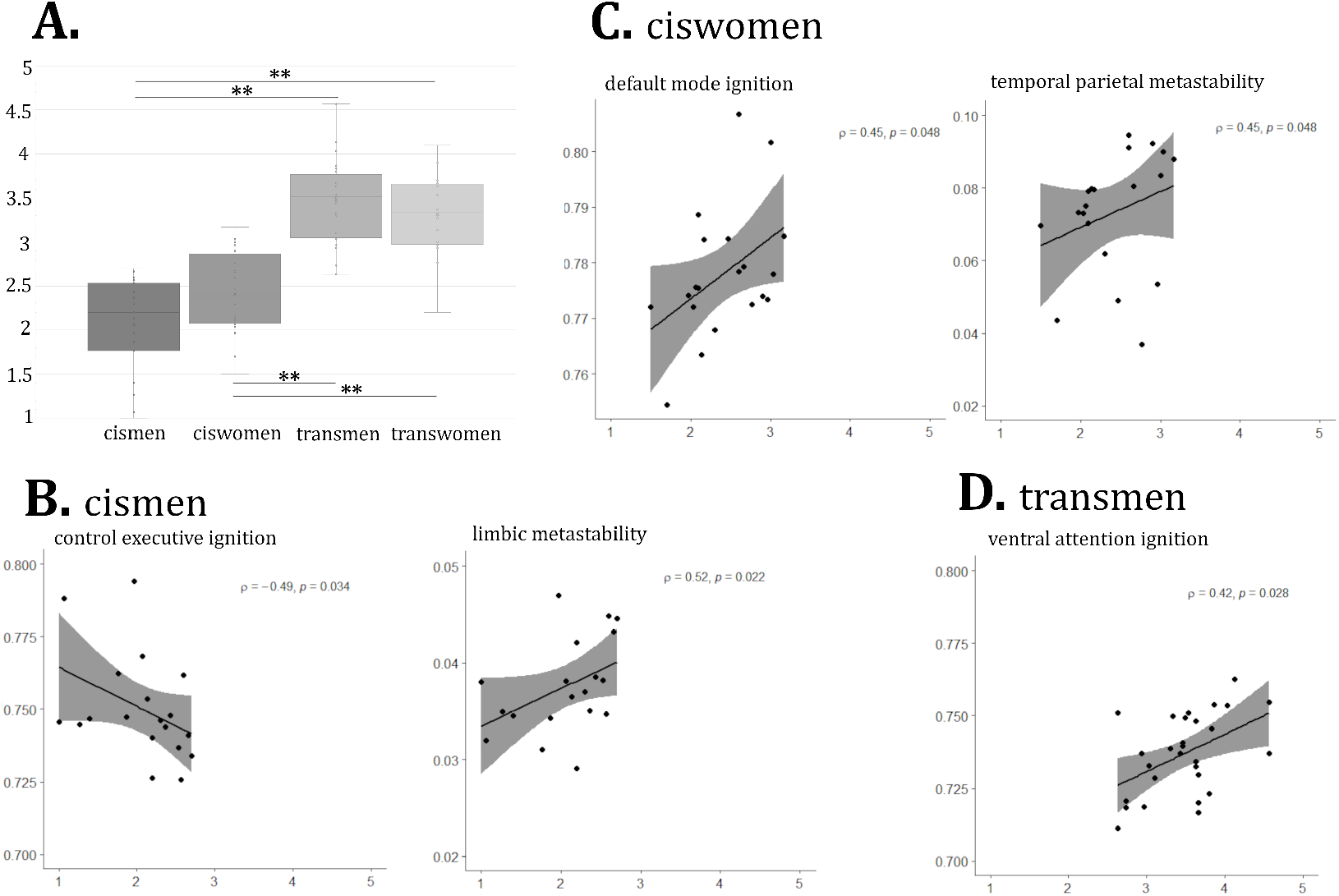
Correlations between network-based ignition and metastability measures and body image satisfaction scores. **(A)** Group comparisons of the body image scale scores. A general linear model with 1,000 permutation testing and Bonferroni was applied. Legend: * *p ≤* 0.05; ** *p* < 0.01. Correlations within **(B)** the cismen group, **(C)** the ciswomen, and **(D)** the transmen. There were no significant correlations within the transwomen group. Correlations are Spearman’s *ρ* and shadowed areas are 95% confidence interval.

Data of mean-ignition and node-metastability matrices per gender group and group comparisons stats are publicly available (https://doi.org/10.6084/m9.figshare.14622564).

## Discussion

For the first time, we characterize the spatio-temporal whole-brain dynamics of binary gender identities, cisgender and transgender. Our findings corroborate the existence of four brain pheno-types (Guillamon et al., 2016; Uribe et al., 2020b) beyond the classical, lately under questioning, conception that the human brain can be split into two configurations, the *male* and the *female* (Legato, 2018). We also demonstrate the need to study whole-brain networks in discriminating gender groups, untangling a complex phenomenon as the experience of a gendered self. To characterize the propagation of information and measure the degree of integration of spontaneously occurring events while at rest, we applied the intrinsic ignition framework (Deco and Kringelbach, 2017; Deco et al., 2017b). Such a framework was very sensitive in detecting functional connectivity differences between young adults grouped by gender. Some of these group functional connectivity differences had been elusive when using stationary functional MRI measurements (Uribe et al., 2020b), and sliding-windows approaches to study brain connectivity states (Uribe et al., 2021). In addition, spatial and temporal brain dynamics measures were uniquely related to the satisfaction towards body parts for cismen, ciswomen, and transmen.

The mean intrinsic ignition is an informative measure of the spatial diversity and broadness of communication across the brain. On the other hand, node-metastability captures the variability over time across the whole brain. Both the spatial and temporal variability that defined each gender group were widespread across the whole brain, including nodes from all functional networks. Likewise, when using a support vector machine algorithm inputting stationary group independent component maps and clinical data as features, four gender groups were obtained predicted by different patterns of brain connectivity (Clemens et al., 2020). In addition, our results stress the importance of using fine-grained connectivity measures to study spatio-temporal oscillations over grand averaged functional connectivity measurements, these latter enabling a more narrowed investigation of differences accountable for gender, and the incongruence felt in the transgender community.

Group differences in both subdivisions of the attentional networks and the executive control were in line with previous findings of functional connectivity differences, both stationary (Uribe et al., 2020a) and dynamic (Uribe et al., 2021). The particular group differences in the dorsal and the ventral subdivisions of the attentional network underline the need to study them separately. More relevant, the spatial broadness of communication of nodes in the default-mode network was higher in ciswomen with respect to cismen and transmen. Increased functional connectivity in default-mode regions has been reported in ciswomen in contrast with cismen (Biswal et al., 2010; Ritchie et al., 2018; de Lacy et al., 2019). Also, in the transgender literature, decreased connectivity in this network regions were found in the transmen group in contrast to cismen (Feusner et al., 2017; Uribe et al., 2020a) and ciswomen (Feusner et al., 2017) while it is not a generalized finding, as other studies had negative reports (Nota et al., 2017; Clemens et al., 2017).

On the other hand, the cismen reported pattern of activation relies on sensory-motor regions (Ritchie et al., 2018). The somatomotor network in the Schaefer parcellation included areas of motor action and sensory inputs of the external world, thus it is the most direct network interacting with our environment. Despite the previous relevance given to such functional network in understanding the own body perception and subsequently explaining the incongruence in transgender people (Manzouri et al., 2017; Burke et al., 2019), the intrinsic ignition framework only differentiated both cisgender groups in terms of temporal variability. Indeed, in our previous work of functional connectivity dynamics (Uribe et al., 2021), a sensorimotor state was found, although no differences between trans and cisgender groups were captured (Uribe et al., 2021).

The spatial and temporal dynamism of the limbic network was greater in the cismen group than in transmen. On the other hand, increased limbic connectivity in transgender individuals has been reported when viewing “ambiguous, androgynous images of themselves morphed toward their gender identity” (Majid et al., 2020). Such findings should be further explored. Different functional MRI measurements do not allow further discussion, and greater integration, broadness of communication, and temporal variability do not necessarily translate to increased averaged connectivity.

The superior parietal cortex has been previously linked to gender differences when comparing cismen with ciswomen and transgender groups, structurally (Zubiaurre-Elorza et al., 2013) and functionally (Uribe et al., 2020a). The choice of the Schaefer parcellation (Schaefer et al., 2018) depicted a high representation of the temporal parietal network in terms of brain dynamics in agreement with temporoparietal junction findings in transmen with respect to cisgender groups (Manzouri et al., 2017). The spatial diversity and broadness of communication of temporal parietal regions were greater in cismen than in the other three gender groups, namely ciswomen, transmen, and transwomen.

The fact that cismen present higher brain dynamism than other gender groups, especially cis-women and transmen, would be in line with previous brain states occupancy where cismen occupied more combinations of connectivity patterns over time than ciswomen (Yaesoubi et al., 2015). Nonetheless, such results have not consistently been replicated, like other brain flexibility measures through brain states using sliding windows reported regional brain dynamism for both cismen and ciswomen differentially (Mao et al., 2017). In addition, the increased spatial and temporal variability of brain oscillations in cismen was not homogeneous for all networks, as in the default-mode network.

Transwomen presented a lateralized predominance in the regions with the highest node-metastability in the left hemisphere. The discussion of such findings is limited due to the scarce literature investigating gender differences in brain dynamism. To the best of our knowledge, previous reports of the gender effects in the lateralization of brain connectivity patterns were mostly comparable between a large sample of cis men and women, with two marginal findings that did not survive false discovery rate correction and were considered a trend-level effect (Agcaoglu et al., 2015; Eliot et al., 2021), and marginal leftwards lateralization in ciswomen only in the inferior frontal cortex (Tomasi and Volkow, 2012). Given these and the small sample of individuals investigated, especially in the transwomen group, our results should be taken carefully.

Finally, the (dis)satisfaction towards the own body parts is not simply associated with a specific network, but differently according to the group, which suggests a different way to understand and accept the body depending on gender. The transmen group image (dissatisfaction) relied on the ventral attentional, i.e., salience network. If one key element in the construction of gender is the perception of our own body (Peelen and Downing, 2007; Burke et al., 2019), the salience network has been highly related to trans and cisgender differences that may explain the gender incongruence (Uribe et al., 2020a, 2021). However, such a landmark is not helpful for the functional correlates of cisgender groups. These differences in the networks correlates could be driven by the fact that the transmen group scores were within the range of the unconformity towards the body parts —4-5 points in the Likert scale of Lindgren and Pauly (1975)—, while cisgender groups would range mainly within the neutral-satisfaction scores (1 to 3 points). Thus, the cismen group’s satisfaction and/or neutrality were positively associated with the limbic network and negatively with the executive control. On the other hand, the network with higher spatial dynamical complexity, i.e., the default-mode, was also associated with body parts satisfaction in ciswomen. Our works provide evidence that the construction of a gendered self, i.e., an inner self with maleness, femaleness, or other variants endowed, is a complex phenomenon characterized by the whole-brain network interplay in terms of spatial and temporal variability that exceed the rather specific correlates of the degree of satisfaction towards the own body.

Some shortcomings should be addressed in future works. First, the aforementioned need to increase the sample size that would add more power to the findings. Collaborative initiatives currently ongoing are trying to overcome such a persistent limitation in the neuroimaging field as for the ENIGMA initiative on transgender health (Mueller et al., 2021). However, it lacks standard acquisition protocols to reduce the variability accounted for sites that may hamper group discrimination. Second, the menstrual cycle of ciswomen and transmen was not accounted as a variable of interest, while there is growing evidence pointing out there are functional connectivity dynamics differences across the menstrual cycle (De Filippi et al., 2021). Likewise, the sexual orientation of all participants was not systematically assessed.

Including minority gender groups when investigating the gender phenomenon in the brain are imperative to understand the complexity of the gender experience. Nonetheless, future studies should include other gender groups, such as nonbinary or other genderqueer identities. Our exploratory work could potentially impact awareness, the development of healthcare guidelines, societal and political evidence-based changes accounting for such heterogeneity, and improving the quality of life while raising visibility that can help fight stigma (Janssen and Voss, 2020).

## Conclusion

First, we propose a gendered brain perspective of spatio-temporal whole-brain communications across networks that characterize four binary gender groups, namely cis- and transmen and women. Second, we stress the need to study the brain as a whole complex system beyond the localizationism paradigm when investigating complex phenomena such as the unique construction of gender. Third, taking advantage of novel brain dynamics techniques to understand network cooperation and the brain’s dynamical complexity, we confirm and expand previous findings relating to the attentional network as the cornerstone of gender differences. However, the default-mode, executive control, limbic, somatomotor, and temporal parietal networks also presented differences in information propagation between cis and transgender identities. Fourth, we propose a unique brain network experience in perceiving satisfaction towards the own body parts according to each gender group. Finally, novel cutting-edge frameworks in studying brain dynamics are needed to untangle the complex and very intimate experience of a gendered self.

## Acknowledgements

CU was supported by the European Union’s Horizon 2020 research and innovation programme under the Marie Sklodowska-Curie grant agreement 888692. AG was supported by the Spanish Ministry of Science and Innovation grant number PGC2018-094919-B-C21. AE was supported by European Union’s Horizon 2020 FET Flagship Human Brain Project (grant agreement number 945539, HBP SGA3). GD was supported by the Spanish Research Project (ref. PID2019-105772GB-I00 AEI FEDER EU), funded by the Spanish Ministry of Science, Innovation and Universities (MCIU), State Research Agency (AEI), and European Regional Development Funds (FEDER); and HBP SGA3 Human Brain Project Specific Grant Agreement 3 (grant agreement no. 945539), funded by the EU H2020 FET Flagship program and SGR Research Support Group support (ref. 2017 SGR 1545), funded by the Catalan Agency for Management of University and Research Grants (AGAUR). EDF was supported by the Doctorate Scholarship FI-2020 from the Catalan Agency for Management of University and Research Grants (AGAUR). The authors are thankful to the participants for their time, without them this work would not be a reality. We are also indebted to the Magnetic Resonance Imaging core facility of the IDIBAPS for technical support, especially to Cesar Garrido and Gema Lasso; and we acknowledge the CERCA Programme/Generalitat de Catalunya.

## Authors contributions

**>Carme Uribe:** Designed research, Analyzed data, Investigation, and Writing - Original draft preparation. **Anira Escrichs:** Designed research, Analyzed data, Software and Writing - Original draft preparation. **Eleonora de Filippi:** Visualization and Writing - Original draft preparation. **Yonatan Sanz-Perl:** Software and Writing - Reviewing and Editing. **Carme Junque:** Super-vision and Writing - Reviewing and Editing. **Esther Gomez-Gil:** Investigation and Writing - Reviewing and Editing. **Morten Kringelbach:** Methodology, Software and Writing - Reviewing and Editing. **Antonio Guillamon:** Funding acquisition and Writing - Reviewing and Editing. **Gustavo Deco:** Methodology, Software and Writing - Reviewing and Editing.

## Data accessibility statement

Imaging raw and processed data used for this study and demographical variables are publicly available in Data Mendeley repositories (Uribe et al., 2020a): https://doi.org/10.17632/hjmfrv6vmg. 2, https://doi.org/10.17632/rw2yhtpj96.3, https://doi.org/10.17632/bgyzz94mz9.3. The data that support the findings of this study are openly available in https://doi.org/10.6084/m9.figshare.14622564.

## References

Agcaoglu O, Miller R, Mayer AR, Hugdahl K, Calhoun VD (2015) Lateralization of resting state networks and relationship to age and gender. Neuroimage 104:310–325.

Allen EA, Damaraju E, Plis SM, Erhardt EB, Eichele T, Calhoun VD (2014) Tracking whole-brain connectivity dynamics in the resting state. Cerebral cortex 24:663–676.

Alonso Martínez S, Marsman JBC, Kringelbach ML, Deco G, ter Horst GJ (2020) Reduced spatiotemporal brain dynamics are associated with increased depressive symptoms after a relationship breakup. NeuroImage Clin. 27:102299.

Biswal BB, Mennes M, Zuo XN, Gohel S, Kelly C, Smith SM, Beckmann CF, Adelstein JS, Buckner RL, Colcombe S, Dogonowski AM, Ernst M, Fair D, Hampson M, Hoptman MJ, Hyde JS, Kiviniemi VJ, Kotter R, Li SJ, Lin CP, Lowe MJ, Mackay C, Madden DJ, Madsen KH, Margulies DS, Mayberg HS, McMahon K, Monk CS, Mostofsky SH, Nagel BJ, Pekar JJ, Peltier SJ, Petersen SE, Riedl V, Rombouts SARB, Rypma B, Schlaggar BL, Schmidt S, Seidler RD, Siegle GJ, Sorg C, Teng GJ, Veijola J, Villringer A, Walter M, Wang L, Weng XC, Whitfield-Gabrieli S, Williamson P, Windischberger C, Zang YF, Zhang HY, Castellanos FX, Milham MP (2010) Toward discovery science of human brain function. Proceedings of the National Academy of Sciences 107:4734–4739.

Bouman WP, Schwend AS, Motmans J, Smiley A, Safer JD, Deutsch MB, Adams NJ, Winter S (2017) Language and trans health.

Burke SM, Majid DSA, Manzouri AH, Moody T, Feusner JD, Savic I (2019) Sex differences in own and other body perception. Human Brain Mapping 40:474–488.

Chen AC, Oathes DJ, Chang C, Bradley T, Zhou ZW, Williams LM, Glover GH, Deisseroth K, Etkin A (2013) Causal interactions between fronto-parietal central executive and default-mode networks in humans. Proceedings of the National Academy of Sciences 110:19944–19949.

Clemens B, Derntl B, Smith E, Junger J, Neulen J, Mingoia G, Schneider F, Abel T, Bzdok D, Habel U (2020) Predictive Pattern Classification Can Distinguish Gender Identity Subtypes from Behavior and Brain Imaging. Cerebral Cortex 30:2755–2765.

Clemens B, Junger J, Pauly K, Neulen J, Neuschaefer-Rube C, Frölich D, Mingoia G, Derntl B, Habel U (2017) Male-to-female gender dysphoria: Gender-specific differences in resting-state networks. Brain and behavior 7:e00691.

David SP, Naudet F, Laude J, Radua J, Fusar-Poli P, Chu I, Stefanick ML, Ioannidis JP (2018) Potential reporting bias in neuroimaging studies of sex differences. Scientific reports 8:1–8.

De Filippi E, Uribe C, Avila-Varela DS, Martínez-Molina N, Pritschet L, Gashaj V, Santander T, Jacobs EG, Kringelbach ML, Sanz Y, Deco G, Escrichs A (2021) The menstrual cycle modulates whole-brain turbulent dynamics. Frontiers in Neuroscience 15:753820.

de Lacy N, McCauley E, Kutz JN, Calhoun VD (2019) Sex-related differences in intrinsic brain dynamism and their neurocognitive correlates. NeuroImage 202:116116.

Deco G, Cruzat J, Cabral J, Tagliazucchi E, Laufs H, Logothetis NK, Kringelbach ML (2019) Awakening: Predicting external stimulation to force transitions between different brain states. Proceedings of the National Academy of Sciences 116:18088–18097.

Deco G, Kringelbach ML (2017) Hierarchy of Information Processing in the Brain: A Novel ‘Intrinsic Ignition’ Framework. Neuron 94:961–968.

Deco G, Kringelbach ML, Jirsa VK, Ritter P (2017a) The dynamics of resting fluctuations in the brain: metastability and its dynamical cortical core. Scientific reports 7:1–14.

Deco G, Tagliazucchi E, Laufs H, Sanjuán A, Kringelbach ML (2017b) Novel intrinsic ignition method measuring local-global integration characterizes wakefulness and deep sleep. eNeuro 4:1–12.

Deco G, Tononi G, Boly M, Kringelbach ML (2015) Rethinking segregation and integration: Contributions of whole-brain modelling. Nature Reviews Neuroscience 16:430–439.

Eliot L, Ahmed A, Khan H, Patel J (2021) Dump the “dimorphism”: Comprehensive synthesis of human brain studies reveals few male-female differences beyond size. Neuroscience & Biobehavioral Reviews 125:667–697.

Escrichs A, Biarnes C, Garre-Olmo J, Fernández-Real JM, Ramos R, Pamplona R, Brugada R, Serena J, Ramió-Torrentà L, Coll-De-Tuero G, Gallart L, Barretina J, Vilanova JC, Mayneris-Perxachs J, Essig M, Figley CR, Pedraza S, Puig J, Deco G (2020) Whole-Brain Dynamics in Aging: Disruptions in Functional Connectivity and the Role of the Rich Club. Cerebral Cortex 31:2466–2481.

Escrichs A, Sanjuán A, Atasoy S, López-González A, Garrido C, Càmara E, Deco G (2019) Characterizing the dynamical complexity underlying meditation. Frontiers in Systems Neuroscience 13:2015–2020.

Feusner JD, Lidström A, Moody TD, Dhejne C, Bookheimer SY, Savic I (2017) Intrinsic network connectivity and own body perception in gender dysphoria. Brain Imaging and Behavior 11:964–976.

Glerean E, Salmi J, Lahnakoski JM, Jääskeläinen IP, Sams M (2012) Functional Magnetic Resonance Imaging Phase Synchronization as a Measure of Dynamic Functional Connectivity. Brain Connectivity 2:91–101.

Guillamon A, Junque C, Gómez-Gil E (2016) A Review of the Status of Brain Structure Research in Transsexualism. Archives of Sexual Behavior 45:1615–1648.

Hansen EC, Battaglia D, Spiegler A, Deco G, Jirsa VK (2015) Functional connectivity dynamics: modeling the switching behavior of the resting state. Neuroimage 105:525–535.

Hyde JS, Bigler RS, Joel D, Tate CC, van Anders SM (2019) The future of sex and gender in psychology: Five challenges to the gender binary. American Psychologist 74:171.

Janssen A, Voss R (2020) Policies Sanctioning Discrimination Against Transgender Patients Flout Scientific Evidence and Threaten Health and Safety. Transgender Health.

Legato MJ (2018) Untangling the gordian knot of human sexuality: What is the biologic basis of variations in sexual phenotype? Gender and the Genome 2:62–67.

Lindgren TW, Pauly IB (1975) A body image scale for evaluating transsexuals. Archives of Sexual Behavior 4:639–656.

Majid DS, Burke SM, Manzouri A, Moody TD, Dhejne C, Feusner JD, Savic I (2020) Neural Systems for Own-body Processing Align with Gender Identity Rather Than Birth-assigned Sex. Cerebral Cortex 30:2897–2909.

Manzouri A, Kosidou K, Savic I (2017) Anatomical and Functional Findings in Female-to-Male Transsexuals: Testing a New Hypothesis. Cerebral cortex (New York, N.Y. : 1991) 27:998–1010.

Mao N, Zheng H, Long Z, Yao L, Wu X (2017) Gender differences in dynamic functional connectivity based on resting-state fMRI In Proceedings of the Annual International Conference of the IEEE Engineering in Medicine and Biology Society, EMBS, Vol. 2017, pp. 2940–2943. Institute of Electrical and Electronics Engineers Inc.

Mayneris-Perxachs J, Arnoriaga-Rodríguez M, Martín M, Burokas A, Blasco G, Coll C, Escrichs A, Biarnés C, Moreno-Navarrete JM, Puig J, Garre J, Ramos R, Pedraza S, Brugada R, Vilanova JC, Serena J, Gich J, Ramió-Torrentà L, Pérez-Brocal V, Moya A, Pamplona R, Sol J, Jové M, Ricart W, Portero-Otin M, Deco G, Maldonado R, Fernandez-Real JM (2021) Microbiota alterations in proline metabolism impact on depression through GABA and ECM homeostasis. Research Square.

Menon V (2011) Large-scale brain networks and psychopathology: A unifying triple network model. Trends in Cognitive Sciences 15:483–506.

Mueller SC, Guillamon A, Zubiaurre-Elorza L, Junque C, Gomez-Gil E, Uribe C, Khorashad BS, Khazai B, Talaei A, Habel U et al. (2021) The neuroanatomy of transgender identity: Mega-analytic findings from the enigma transgender persons working group. The Journal of Sexual Medicine.

Nota NM, Burke SM, den Heijer M, Soleman RS, Lambalk CB, Cohen-Kettenis PT, Veltman DJ, Kreukels BP (2017) Brain sexual differentiation and effects of cross-sex hormone therapy in transpeople: A resting-state functional magnetic resonance study. Neurophysiologie Clinique 47:361–370.

Padilla N, Saenger VM, Van Hartevelt TJ, Fernandes HM, Lennartsson F, Andersson JL, Kringelbach M, Deco G, °Aden U (2020) Breakdown of Whole-brain Dynamics in Preterm-born Children. Cerebral Cortex 30:1159–1170.

Peelen MV, Downing PE (2007) The neural basis of visual body perception. Nature reviews neuroscience 8:636–648.

Polderman TJ, Kreukels BP, Irwig MS, Beach L, Chan YM, Derks EM, Esteva I, Ehrenfeld J, Heijer MD, Posthuma D, Raynor L, Tishelman A, Davis LK (2018) The Biological Contributions to Gender Identity and Gender Diversity: Bringing Data to the Table. Behavior Genetics 48:95–108.

Ritchie SJ, Cox SR, Shen X, Lombardo MV, Reus LM, Alloza C, Harris MA, Alderson HL, Hunter S, Neilson E, Liewald DCM, Auyeung B, Whalley HC, Lawrie SM, Gale CR, Bastin ME, Mcintosh AM, Deary IJ (2018) Sex Differences in the Adult Human Brain: Evidence from 5216 UK Biobank Participants. Cerebral Cortex 28:2959–2975.

Schaefer A, Kong R, Gordon EM, Laumann TO, Zuo XN, Holmes AJ, Eickhoff SB, Yeo BTT (2018) Local-Global Parcellation of the Human Cerebral Cortex from Intrinsic Functional Connectivity MRI. Cerebral Cortex 28:3095–3114.

Selvaggi G, Bellringer J (2011) Gender reassignment surgery: an overview. Nature Reviews Urology 8:274.

Tagliazucchi E, Balenzuela P, Fraiman D, Chialvo DR (2012) Criticality in large-scale brain fmri dynamics unveiled by a novel point process analysis. Frontiers in Physiology 3 FEB.

Thomas Yeo BT, Krienen FM, Sepulcre J, Sabuncu MR, Lashkari D, Hollinshead M, Roffman JL, Smoller JW, Zöllei L, Polimeni JR, Fisch B, Liu H, Buckner RL (2011) The organization of the human cerebral cortex estimated by intrinsic functional connectivity. Journal of Neurophysiology 106:1125–1165.

Tomasi D, Volkow ND (2012) Laterality patterns of brain functional connectivity: gender effects. Cerebral Cortex 22:1455–1462.

Uribe C, Junque C, Gómez-Gil E, Abos A, Mueller SC, Guillamon A (2020a) Data for functional MRI connectivity in transgender people with gender incongruence and cisgender individuals. Data in Brief 31:105691.

Uribe C, Junque C, Gómez-Gil E, Abos A, Mueller SCS, Guillamon A (2020b) Brain network interactions in transgender individuals with gender incongruence. NeuroImage 211:116613.

Uribe C, Junque C, Gómez-Gil E, Díez-Cirarda M, Guillamon A (2021) Brain connectivity dynamics in cisgender and transmen people with gender incongruence before gender affirmative hormone treatment. Scientific Reports 11:1–11.

Yaesoubi M, Miller RL, Calhoun VD (2015) Mutually temporally independent connectivity patterns: A new framework to study the dynamics of brain connectivity at rest with application to explain group difference based on gender. NeuroImage 107:85–94.

Zubiaurre-Elorza L, Junque C, Gómez-Gil E, Segovia S, Carrillo B, Rametti G, Guillamon A (2013) Cortical thickness in untreated transsexuals. Cerebral cortex (New York, N.Y. : 1991) 23:2855–2862.

